# A cortical pathway modulates sensory input into the olfactory striatum

**DOI:** 10.1101/235291

**Authors:** Kate A. White, Yun-Feng Zhang, Zhijian Zhang, Janardhan P. Bhattarai, Andrew H. Moberly, Estelle in ‘t Zandt, Huijie Mi, Xianglian Jia, Marc V. Fuccillo, Fuqiang Xu, Minghong Ma, Daniel W. Wesson

## Abstract

Sensory cortices process stimuli in manners essential for perception. The piriform ‘primary’ olfactory cortex (PCX) extends dense association fibers into the ventral striatum’s olfactory tubercle (OT), yet the function of this cortico-striatal pathway is unknown. We optically stimulated channelrhodopsin-transduced PCX glutamatergic neurons or their association fibers while recording OT neural activity in mice performing an olfactory task. Activation of PCX neurons or their association fibers within the OT controlled the firing of some OT neurons and bidirectionally modulated odor coding dependent upon the neuron’s intrinsic odor responsivity. Further, patch clamp recordings and retroviral tracing from D1 and D2 dopamine receptor-expressing OT medium spiny neurons revealed this input can be monosynaptic and that both cell types receive most of their input from a specific spatial zone localized within the ventro-caudal PCX. These results demonstrate that the PCX functionally accesses the direct and indirect pathways of the basal ganglia within the OT.

## Introduction

How does the brain distribute sensory information to enable the successful encoding of stimuli? In nearly all mammalian sensory systems, information from the environment is transduced by peripheral sensory receptors and relayed into the thalamus, where this input is processed and then distributed into cortical structures that encode stimulus attributes. For example, in the visual system, photoreceptor cells send input into the thalamus for processing, after which the information is distributed into cortical structures which each participate in the encoding of distinct visual features (Kravitz et al., 2013). This general organizational scheme (receptor neurons → thalamus → cortical structures) is utilized by all sensory systems but the olfactory system. This leads to the notion that within the olfactory system, it is the interaction of olfactory cortical structures which is integral for the generation of an odor percept (Haberly, 2001).

In the mammalian olfactory system, odors are transduced by olfactory sensory neurons in the nasal epithelium which extend their axons into the olfactory bulb wherein they coalesce to form glomeruli (Shepherd et al., 2004; Wilson and Mainen, 2006). From here, mitral and tufted cells which are the principal neurons of the olfactory bulb, relay this odor information into secondary olfactory structures, including the piriform cortex (PCX), olfactory tubercle (OT), anterior olfactory nucleus, and cortical amygdala, among others (Haberly, 2001). Notably, in addition to extensive bulbar connections, including some which are reciprocal, these cortical structures are also heavily interconnected (for review see (Giessel and Datta, 2014; Haberly, 2001; Neville and Haberly, 2004)). Thus, across the olfactory system, there is diffuse connectivity that allows for the potential to transform odor information.

The PCX, often referred to as the ‘primary’ olfactory cortex, extends massive numbers of glutamatergic association fibers throughout the brain which innervate other olfactory structures (Haberly, 2001; Neville and Haberly, 2004; Schwob and Price, 1984a; Shipley and Ennis, 1996). PCX neural ensembles precisely encode certain odor features (Bolding and Franks, 2017; Poo and Isaacson, 2009; Rennaker et al., 2007; Stettler and Axel, 2009; Wilson, 2000a, 2000b), thereby providing the possibility that the association fiber network serves to relay odor information among brain structures. Additionally, local association fibers within the PCX shape the encoding of odors via plastic actions and recurrent circuitry (Barkai et al., 1994; Franks et al., 2011; Hasselmo et al., 1990; Large et al., 2016; Linster et al., 2009; Suzuki and Bekkers, 2006, 2011). Do PCX association fibers shape the representation of odors in interconnected olfactory structures? Here we tested this question by interrogating the influence of PCX association fiber input on the OT.

The OT receives direct input from the olfactory bulb (Haberly and Price, 1977; Scott et al., 1980), and like the PCX, it also participates in odor coding, such as encoding odor identity and intensity (Payton et al., 2012; Xia et al., 2015) and assigning an odor with valence (Gadziola et al., 2015). However, the OT is the only olfactory structure which is also a component of the ventral striatum (Heimer et al., 1982). This characteristic is postulated to afford the OT with the capacity to shape sensory-directed motivated behaviors (Ikemoto, 2007; Wesson and Wilson, 2011). As is the case with other striatal structures, the principal class of neurons in the OT are medium spiny neurons (MSNs) (Meredith, 1999). These MSNs express either D1- or D2-type dopamine receptors, with striatopallidal MSNs (those mostly projecting to the ventral pallidum/globus pallidus) expressing D2 receptors and striatonigral MSNs (innervating substantia nigra or ventral tegmental area) expressing D1 receptors (Gerfen et al., 1990; Zhang et al., 2017) (for review (Smith et al., 2013; Tian et al., 2010)). Thus, the OT is preferentially situated, as the olfactory striatum, to send sensory information into downstream basal ganglia structures via the direct (D1) and indirect (D2) pathways, extending projections into both midbrain and other striatal structures (Heimer and Wilson, 1975; Wesson and Wilson, 2011).

We predicted that the PCX shapes odor processing within the OT by means of the PCX association fiber network. PCX association fibers innervate all layers of the OT (Johnson et al., 2000; Schwob and Price, 1984a, 1984b), and stimulation of the PCX *ex vivo* elicits postsynaptic potentials in the OT (Carriero et al., 2009). *In vivo* recordings also suggest that network oscillations in the ventral striatum originate in the PCX (Carmichael et al., 2017). It remains unknown whether, and if so how, PCX inputs modulate olfactory processing in the OT. Further, the spatial population of neurons within the PCX that innervate the OT, and the OT neuron types receiving this PCX input, are both unresolved.

To address these voids, we recorded the activity of OT neurons in behaving mice while optically stimulating channelrhodopsin (ChR2)-expressing principal neurons in the PCX and/or their association fibers in the OT while mice performed an olfactory task. We then employed patch-clamp recordings and viral tracing methods to determine the connectivity of PCX neurons upon OT D1 and D2 MSNs. Our work uncovers that the PCX controls the number of odor responsive OT neurons and the direction of their response. Moreover, OT D1 and D2 MSNs receive monosynaptic inputs from neurons which are topographically organized to be largely within the ventro-caudal anterior PCX.

## Results

### Viral strategy for the optogenetic control of PCX principal neurons

In order to target PCX neurons and their association fibers innervating the OT for later optogenetic manipulation, we injected an AAV viral vector designed to express ChR2 and a reporter fluorophore (mCherry) under control of the calcium/calmodulin-dependent protein kinase iiα (CaMKiiα) promoter, focally, in the anterior PCX (AAV5.CaMKiiα.hChR2(E123T/T159C).p2A.mCherry.WPRE or AAV5.CaMKiiα.hChR2(H134R).mCherry; **Fig 1A**). This approach led to infection of PCX neurons (**Fig 1B-C**). Anti-NeuN immunohistochemistry on tissue from a subset of mice not used in recordings was utilized to label neurons and to aid in quantification of viral infection efficiency. 59.1% of NeuN-labeled neurons within a set PCX region of interest were also mCherry+ (inter-animal range 51.5 – 66.2% ± 5.6% s.d.; **Fig 1C**, see Methods). From all colabeled neurons, the greatest numbers were found in layer ii (78.9%, mean = 101.1 neurons), followed by layers iii and i, respectively (iii: 16.9%, mean = 21.7 neurons; i: 4.2%, mean = 5.5 neurons; *F*(*2,7*)=206.47, *p*<0.0001; **Fig 1D**). Importantly, layer ii contains the densest collection of projection neurons (superficial pyramidal neurons and semilunar cells) extending fibers into the OT (Bekkers and Suzuki, 2013; Haberly, 2001; Mazo et al., 2017; Neville and Haberly, 2004; Shipley and Ennis, 1996). While this AAV is, by design, specific for transducing excitatory neurons, we nevertheless confirmed that the neurons infected with AAV and thus expressing ChR2 are indeed glutamatergic by performing anti-VGlut1 immunohistochemistry. Colabeling of mCherry and anti-Vglut1 was observed in these cases which further confirms cellular specificity of the AAV infection (**Fig S1D**).

**Figure 1.**
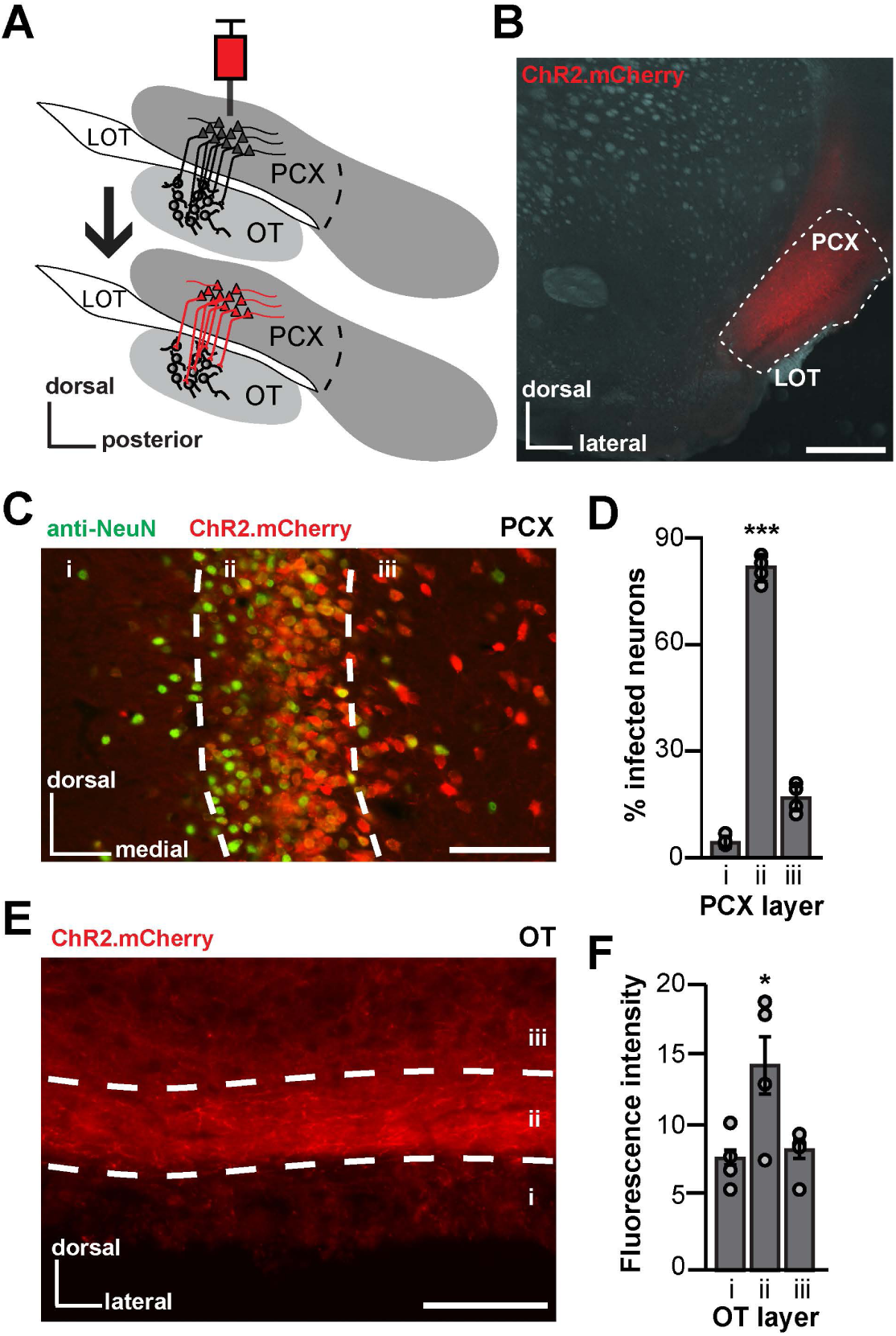
Viral strategy for the optogenetic control of PCX principal neurons. **A.** Schematic of the AAV injection procedure into the anterior PCX that was used in order to express ChR2 in PCX neurons and association fibers. LOT = lateral olfactory tract, PCX = piriform cortex, OT = olfactory tubercle. Following injection of AAV into the PCX (top) 23 weeks of time was allowed for viral transduction, following which mice were used for physiological experiments (bottom). **B**. Representative image of a PCX AAV injection (AAV5.CaMKiia.hChR2(H134R).mCherry) restricted into specifically the anterior PCX (white dotted outline). Scale bar = 500 μm. **C**. Representative image of PCX neurons (anti-NeuN, marker for neuronal nuclei) transduced with the viral vector AAV5.CaMKiiα.hChR2.E123T.T159C.p2A.mCherry.WPRE across PCX layers i-iii. Scale bar = 100 μm. D. Quantification of AAV transduction in the PCX across animals (*n*=4) as a percentage of mCherry+ cells compared to NeuN+ cells in each PCX cell layer. \*\*\**p*<0.001. **E**. Image of mCherry+ association fibers within the OT which originated from the PCX. Scale bar = 100μm. **F**. The average fluorescence intensity of PCX association fibers in each OT layer across animals (*n*=4) as a function of background fluorescence. See **Fig S1** for additional histological analyses.

Using this viral approach, we also observed mCherry+ PCX association fibers innervating the OT **(Fig 1E).** As anticipated based upon previous tracing studies (Schwob and Price, 1984a, 1984b), these fibers were observed in all cell layers of the OT. To quantify the density of these association fibers, we extracted the fluorescence intensity across OT cell layers from 4 mice. Fluorescence intensity was greatest in layer ii, followed by layers iii and i (*F*(*2,9*)=4.63, *p*=0.04) (**Fig 1F**). Thus, this AAV-based approach allows for the targeting of ChR2 into PCX glutamatergic neurons and their association fibers, including those innervating the OT. We utilized this same viral approach in two *in vivo* and one *in vitro* opto-physiological paradigms as described next.

### Activation of PCX neurons enhances OT activity

We next sought to address whether the PCX influences the activity of OT neurons *in vivo.* Three weeks after viral injection as described above, mice were implanted with an optical fiber in the PCX for light stimulation of ChR2-transduced neurons (**Fig S1B**) and an electrode array in the OT to record single unit neural activity (**Fig 2A**). In the same surgery, mice were implanted with a head bar for subsequent head-fixation. Following several days of recovery, the mice were water deprived and trained to perform a fixed-interval olfactory task (**Fig S2**) in which head-fixed mice received one of four odors for 2 seconds either with or without simultaneous PCX stimulation consisting of a blue light stimulus train. Mice also received pseudorandom trials of light alone. Odors were presented during the inter-trial interval prior to a window of reward availability, where mice licked to receive a fluid reinforcer. In this design, we ensured the mice were indeed engaged throughout recording sessions by monitoring the occurrence of licking during successive reward windows (**Table S1**). This fixed-interval reinforcement paradigm, which provides reinforcement between all odors in a manner not closely linked in time with odor delivery ensures all odors are of similar valence to the mice.

**Figure 2.**
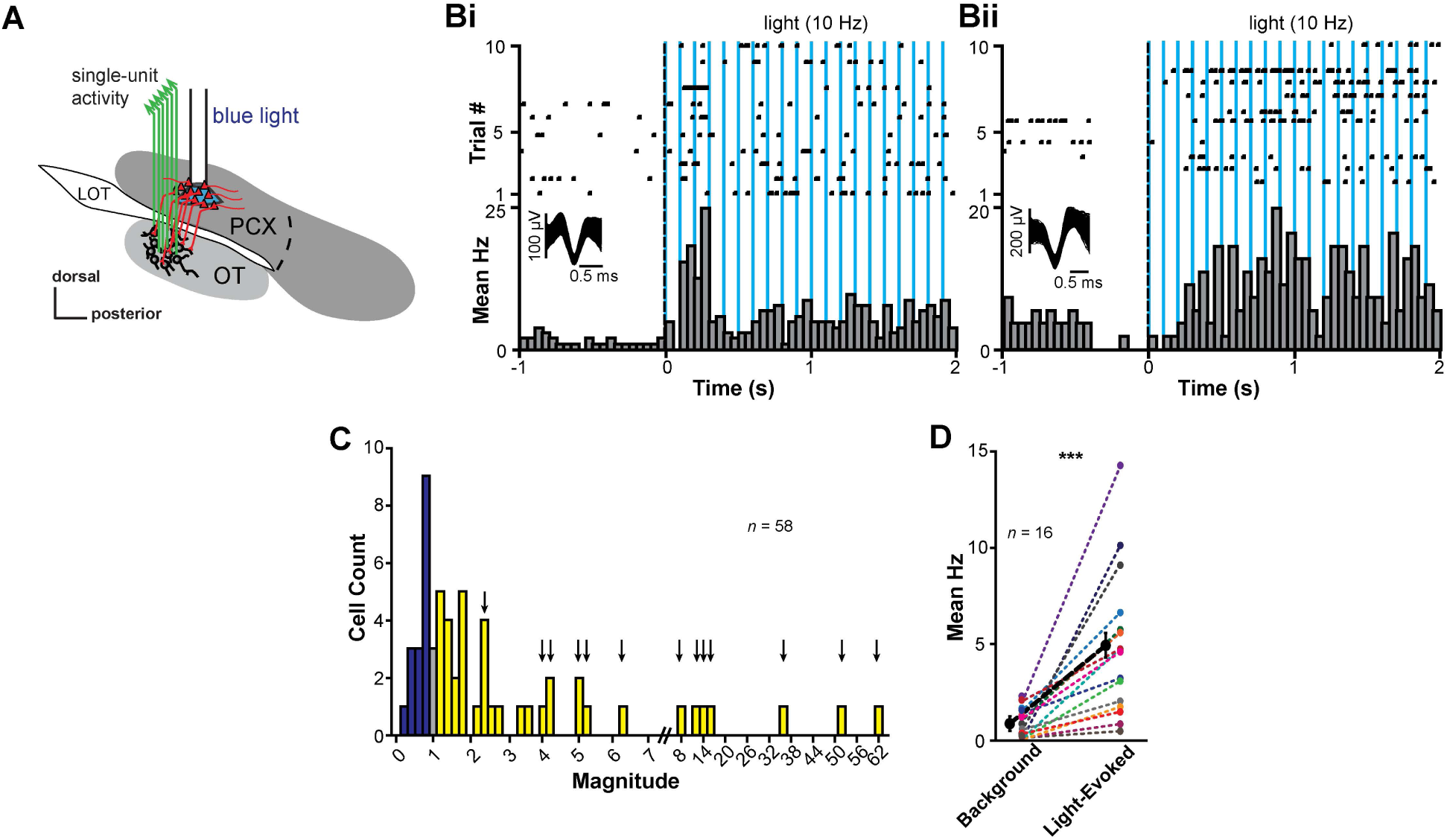
Modulation of OT activity by PCX neuron activation. **A**. Schematic of the location of the optical fiber in PCX for light stimulation and the fixed electrode array in the OT for extracellular recordings. **B**. Examples of significant light-evoked activity of two OT single units by stimulation of the PCX. Each raster plot and histogram represent a single unit’s response to light stimulation across 10 trials. Average waveforms for each unit are represented within the histogram. Responses to PCX light stimulation evokes unique temporal excitation, with activity either transient (i) or sustained (ii). **C**. Distribution of the average light-evoked response across a population of recorded units. A majority displayed excitation to PCX light stimulation (yellow), while some were suppressed (purple) and a small number remained unchanged (gray). Arrows indicate bars where units contributed to the plot in **D**. Double-line break in x axis denotes change in magnitude binning (from 0.2 to 2) to represent sizable changes in magnitude upon light stimulation. **D**. Distribution of the change from average background activity to light-evoked activity. Of 16 significantly light-responsive units, all units increased their firing rates, demonstrated in the line graph. As a population, these units were significantly excited from background activity. The mean of this data is denoted by the bold black dashed line in **D**. See **Fig S3** for additional light-evoked activity quantification across preparations, and activity from control animals.

Through both *post-mortem* analyses and behavioral scoring, we focused our analyses on mice which met the following four criteria: 1) AAV expression verified to be restricted within the PCX, 2) electrode arrays confirmed within the OT, 3) fiber tips localized within the PCX, and 4) criterion-level behavioral performance (**Fig S1Ai; Table S1**). Among these mice (n=8), 6 had clear multi-unit activity from which single units were sorted. The majority of OT neurons display low background firing rates (Gadziola et al., 2015; Xia et al., 2015), providing the possibility that even subtle changes in firing may shape network function. Indeed, the mean background firing rate of the OT units sampled in this experiment was 2.3 ± 1.2 Hz (inter-unit range: 0.0-5.5 Hz; *n*=58 units; **Fig S3Ai**). We asked whether, and if so how, PCX neuron stimulation may modulate this low OT neuron firing rate. To do this, for every unit we compared 2 seconds of averaged background activity to 2 seconds of averaged light-evoked activity and quantified the magnitude and direction of change upon light stimulation (11-21 trials/unit).

We found that light stimulation of ChR2-transduced CaMKiia neurons within the PCX elicits changes in the firing rates of OT units (**Fig 2B-D**). This modulation may occur in the form of brief phasic increases in firing, or those in which the firing is somewhat sustained throughout the light pulses (**Figs 2Bi-Bii**). 27.6% of OT units were significantly modulated by PCX stimulation (16/58 units; observed in 3 out of 6 mice; *p*<0.05, paired *t*-tests; **Fig 2C**). The impact of PCX activation was significant at the population level across all of these modulated units (*t*(*15*)=-4.79, *p*=0.00012; **Fig 2D**). This effect was exclusively due to excitation among these OT units, with all units increasing their firing rate upon light stimulation (**Fig 2C-D**). Some units showed dramatic increases in firing compared to their low background firing rates (*e.g.,* from ~2Hz to near 15Hz; **Fig 2D**). Importantly, the fact that we observed ChR2-dependent modulation of OT units in only 50% of the mice with detectable single units highlights that this effect is not due to nonspecific influences of light stimulation on the neurons (artifact) or the animals (*e.g.,* arousal). Further supporting this, light-evoked responses were not observed in two separate mice injected with AAV5.CaMKiia.mCherry (*n*=28 units; *p*≥0.195, paired *t*-tests; comparing background and light stimulation periods for each unit; **Fig S3E**). Thus, activation of glutamatergic PCX neurons enhances OT neural activity.

### Activation of PCX association fibers within the OT modulates OT activity

Is the PCX capable of influencing OT activity directly through its association fiber system? To test this, we adapted a new *in vivo* preparation wherein we used the same viral approach described above along with an optetrode to directly stimulate PCX association fibers specifically within the OT while simultaneously recording OT unit activity (**Fig 3A**). This approach allowed us to determine the direct contributions of the PCX upon the OT, while minimizing potential bi- or multi-synaptic influences from other interconnected brain structures which may serve to relay activity from the PCX into the OT. Through both *postmortem* analyses and behavioral scoring, we focused our analyses on 8 mice which met the following three criteria: 1) AAV expression verified to be restricted within the PCX, 2) optetrode arrays confirmed within the OT, and 3) criterion-level behavioral performance (**Table S1**). Among these mice, we extracted 203 single units. Corroborating the previous preparation, the spontaneous firing rates of these OT units were also low, at 2.1 Hz ± 3.2 Hz (inter-unit range: 0-26.6 Hz; **Fig S3Aii**). Supporting our hypothesis that PCX may influence the OT monosynaptically through its association fiber network, we also observed significant light-evoked responses in this preparation. Overall, the population-level impact of this modulation was less than that we observed upon local PCX activation (*i.e.* **Fig 2D**; χ^2^(*1*)=13.97, p=0.002, comparing modulated units in both experiments). Nevertheless, light-evoked responses were observed in 5 out of the 8 mice tested (**Fig S3B-D**). Among these mice, 5.1% of all units were significantly modulated (8/156 units; *p*<0.05, paired *t*-test; **Fig S3B-D**). Two of these units displayed significant excitation upon association fiber stimulation (*p*<0.05 paired *t*-tests). In 6 units, association fiber stimulation resulted in suppressed background firing rates (*p*<0.05, paired *t*-tests) which was significant at the population-level (*t*(*5*)=2.96, *p*=0.032; **Fig S3C**). Further, evoked activity in some units was seemingly well-coupled with each light pulse within the stimulus train (**Fig S3Dii**). Thus, activation of PCX fibers within the OT itself is capable of modulating the activity of some OT neurons.

**Figure 3.**
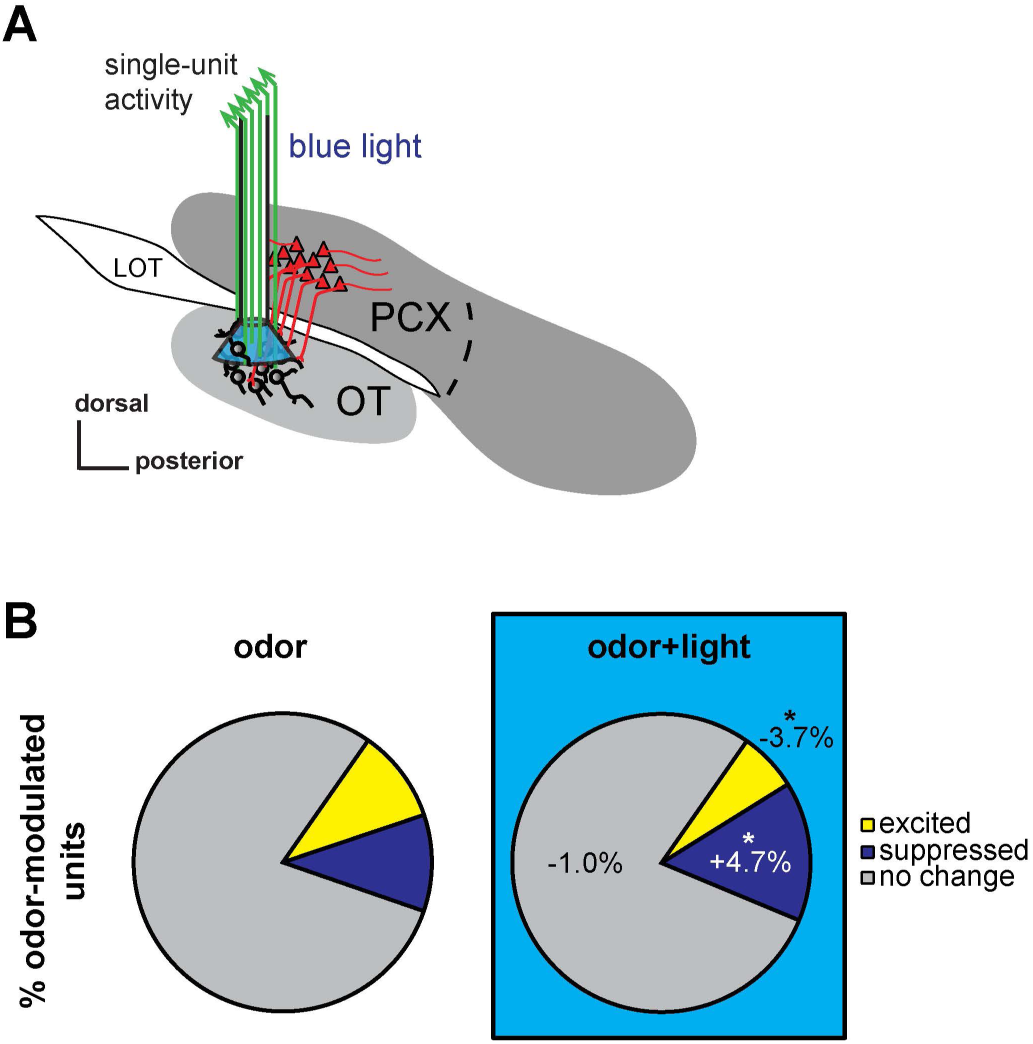
PCX association fiber activation biases OT odor representation. **A.** Schematic depicting the location of the optetrode assembly in the OT to record OT extracellular activity and light stimulate PCX association fiber terminals within the OT. **B**. Population change of odor-responsive OT neurons during PCX light stimulation. Pie charts representing the percent of neurons within the population that were odor-excited (yellow), odor-suppressed (purple), or remain unchanged (gray) with (right) or without (left) PCX light stimulation. \**p*<0.05. See **Fig S2** for olfactory task description.

### Activation of PCX association fibers bidirectionally modulates the representation of odors in the OT

Does PCX input influence the representation of odors in the OT? We reasoned that driving PCX input onto OT neurons, like that which we found excited neurons during either activation of glutamatergic PCX neurons or their association fibers in the OT, would modulate OT neuron firing rates, thereby altering the representation of odors. As described earlier, mice were presented with pseudorandom trials of four different odors, trials of light alone, or trials of light simultaneously with one of the four odors (odor+light) (**Fig S2**). We quantified the number of neurons responsive to each of the odors presented with or without activation of PCX association fibers *via* the optetrode. Across the population of 203 single units, 47.8% (*n*=97; *p*<0.05 in *t*-tests from background to odor-evoked activity) were significantly modulated by at least one of the odors presented. From the activity of these 203 single units, we extracted 812 cell-odor pairs to examine the proportion of units responding to odor compared with simultaneous odor and light. As expected, some cell-odor pairs responded to odors in the form of odor-evoked suppression, whereas others responded by excitation of their firing rates (**Fig 3B**). Not surprisingly, many OT neurons were not modulated by odor (Gadziola et al., 2015). Upon association fiber activation, the proportion of cell-odor pairs significantly excited by odor decreased 3.7% (82 to 52 cell-odor pairs; χ^2^(*1*)=5.77, *p*=0.016), while the proportion of units significantly suppressed by odor increased 4.7% (85 to 123 cell-odor pairs; χ^2^(*1*)=5.80, *p*=0.016). This in some instances was due to light-modulation of units not previously modulated by odor in the absence of light. Thus, activating PCX association fibers within the OT changes the proportion of units representing odors.

Our results demonstrating that PCX association fiber activation alters the proportion of odor-modulated OT units next led us to ask if this activation changes a neuron’s odor-evoked firing rate based upon the way that the neuron responds to odors intrinsically (*viz.,* in the absence of light). To address this question, we quantified the differences in odor responses across individual OT cell-odor pairs. Of the 812 cell-odor pairs, 19.6% (*n*=159) were significantly responsive to at least one of the four odors presented when compared to each cell-odor pair’s average background firing rate (*p*<0.05, within cell-odor pair *t*-tests). 52.2% (*n*=83; **Fig 4Ai-iii**) of these modulated cell-odor pairs were excited by odor presentation, whereas 47.8% displayed odor-evoked suppression of firing (*n*=76; **Fig 4Bi-iii**). To determine the direction of change within each population, we compared each cell-odor pair’s odor-evoked response to that elicited by simultaneous odor and light. Dependent upon whether the cell-odor pair was odor-excited or odor-suppressed, OT units displayed a bidirectional response to simultaneous PCX fiber activation. Those units excited by odors were, as a population, less-excited by the same odors in the context of PCX fiber activation (*t*(*82*)= 5.13, *p*<0.0001, **Fig 4Aii**). This effect was most prominent in cell-odor pairs that encoded odors (in absence of light) with low firing rates (**Fig 4Aiii**). Conversely, odor-suppressed units displayed significant excitation during association fiber activation (*t*(*75*)= −5.74, *p*<0.0001, **Fig 4Bii**). The change in the distribution of cell-odor pairs encoding odor significantly shifted, suggesting that cell-odor pairs with lower odor-evoked firing rates were most greatly modulated by the occurrence of odor and light together (*D*=0.50, *p*=0.023, Komolgorov-Smirnoff test; **Fig 4Biii**). Importantly, a similar bidirectional modulation of OT odor coding was also observed in our previous preparation wherein we stimulated the PCX directly (**Fig S4**). Together, these results demonstrate that PCX association fiber activation biases OT units to encode odors in an opposing manner than they do intrinsically.

**Figure 4.**
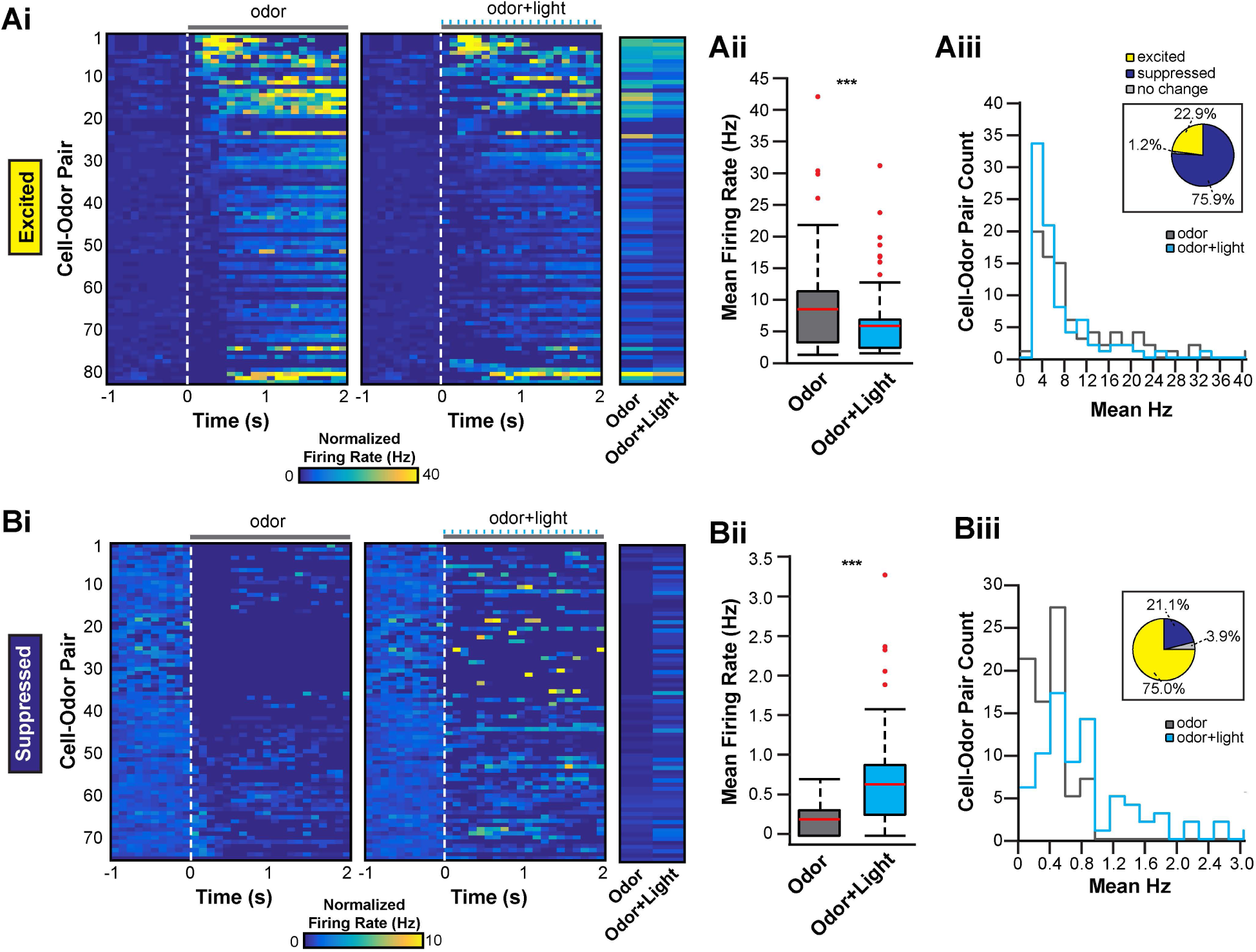
The PCX bidirectionally modulates OT odor-responsive neurons. **A-B.** Modulation of cell-odor pair response from odor alone to odor+light stimulation together. A. represents cell-odor pairs significantly excited by odor, while B. represents cell-odor pairs significantly suppressed by odor. Panels i-iii use the same conventions for both **A**. and **B. i.** 2D histograms representing the change in firing rate from 1 second of background activity to 2 seconds of either odor presentation (left) or odor+light presentation (right). Each row is one unique cell-odor pair and is the same for both left and right 2D histograms. Each column is a 100 ms average firing rate bin for that cell-odor pair. End 2D histogram shows the average evoked response (either odor or odor+light) for each cell-odor pair. **ii.** Boxplot displaying the change in average firing rate across the population for odor alone (gray box) v. odor+light together (blue box). Both plots are significant using a paired *t*-test comparing odor v. odor+light. *p*<0.001. Red line designates the mean, the box designates the 50^th^ percentile range, and the whiskers designate the 25^th^ and 75^th^ percentiles. Red dots are outliers. **iii.** Shows distribution of change from odor alone (gray line) to odor+light together (blue line) from mean firing rate evoked in each case. Pie chart above shows the percent change from odor alone. See **Fig S4** for bidirectional modulation of units during local PCX neuron stimulation.

### PCX principal neurons synapse with, and evoke monosynaptic responses within, OT D1-and D2-type MSNs

The predominant cell type in the OT are MSNs which can be divided into those expressing either the D1 or D2 receptor. Given the uniquely important signaling pathways and downstream connectivity of these different neuron types (Smith et al., 2013; Tian et al., 2010), we next addressed whether PCX input is preferentially directed towards D1 or D2 MSNs. To answer this question, we used an *in vitro* paradigm in combination with the same viral approach we employed *in vivo* – however, the injections were performed in D1-tdTomato/D2-EGFP double-transgenic mice to allow for genetically-guided identification of D1 or D2 neurons, respectively (**Fig S5A**). Three to six weeks after AAV injection, brain slices were collected and whole-cell patch clamp recordings performed on OT D1 and D2 MSNs (**Fig 5A**). Consistent with previous electrophysiological characterizations of striatal MSNs (Cepeda et al., 2008), D1 and D2 MSNs had low input resistances (D1 vs D2: 161.7 ± 7.6 vs. 159.1 ± 9.3 MΩ, *n* = 20 cells in each group; *t*(*38*)=0.21, *p*=0.83). However, D1 MSNs were less excitable compared to D2 MSNs as evidenced by their display of higher firing thresholds and lower firing frequencies upon current injection (Kreitzer and Malenka, 2007) (**Fig S5B-C**).

**Figure 5.**
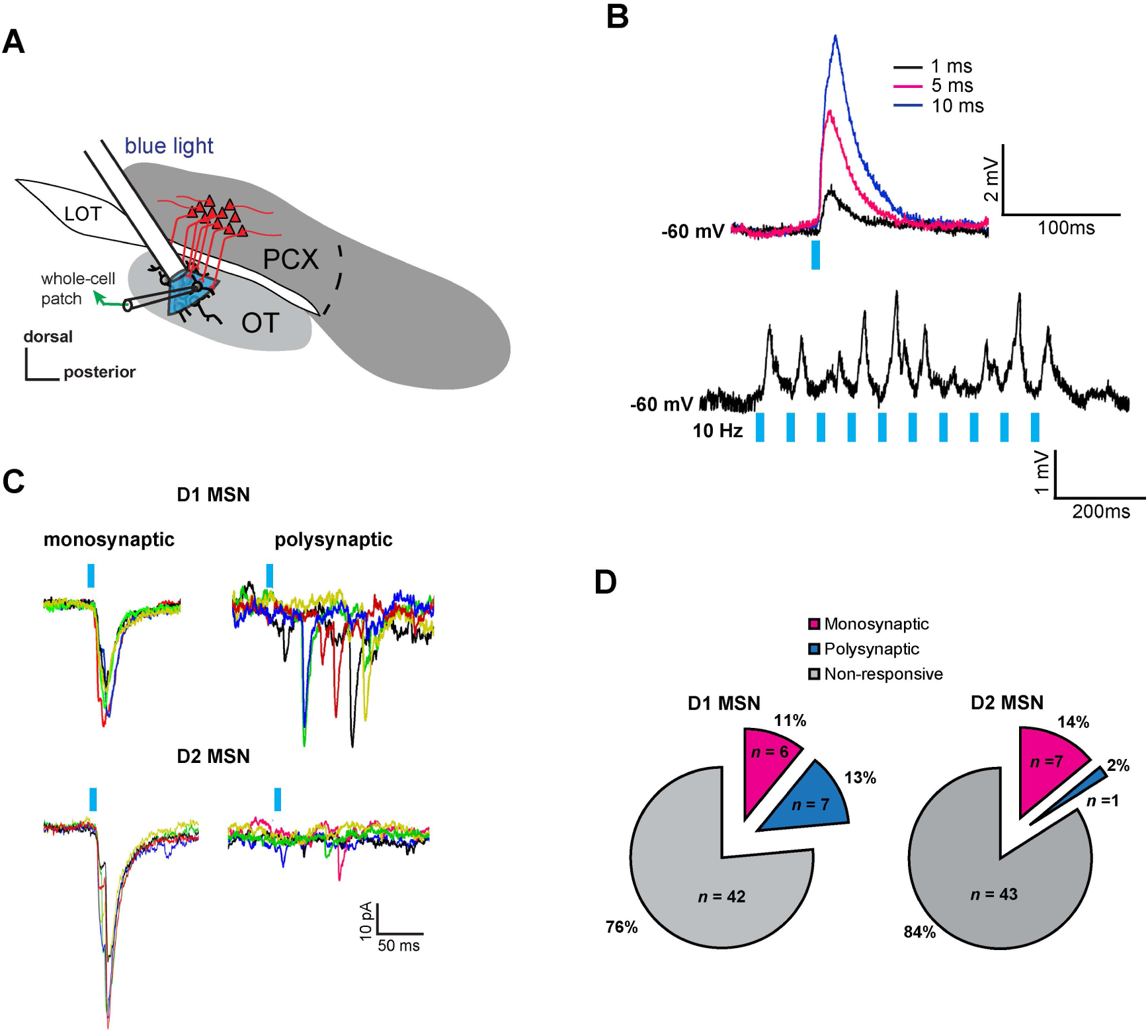
Both OT D1 and D2 MSNs receive monosynaptic and polysynaptic input from the PCX. **A.** Schematic of the whole-cell patch clamp recording of OT D1 or D2 MSNs and stimulation of PCX association fibers. **B**. A D1 MSN in the OT showed light induced EPSPs to different pulse lengths (upper panel) and to a train of 10 Hz stimuli (lower panel) under current clamp mode. **C**. Light-evoked mono- or poly-synaptic EPSCs in D1 and D2 MSNs under voltage clamp mode. The holding potential was −60 mV. **D**. Summary of D1 and D2 cells which displayed light-evoked synaptic responses, organized by whether the evoked response was defined as mono- or polysynaptic (See Methods). See **Fig S5** for additional patch clamp recordings and analysis.

As expected, ChR2-expressing principal neurons in the PCX showed instant (<1 msec latency) and largely repeatable responses to blue light stimulation (**Fig S5D-E**). None of the 106 OT neurons recorded from (55 D1 and 51 D2 MSNs) showed instant light-evoked responses, indicating that they were not directly infected by the virus. In a fraction of D1 MSNs in the OT (**Fig 5B**), light pulses evoked excitatory postsynaptic potentials (EPSPs) under current clamp mode. The evoked responses increased with the pulse duration and typically were able to follow 10 Hz stimulations. To differentiate OT neurons that receive mono- or polysynaptic connections, we recorded light-evoked excitatory postsynaptic currents (EPSCs) under voltage clamp mode and measured the response latency during repeated stimulations (**Fig 5C**). An OT neuron was considered to receive monosynaptic inputs from the PCX if the response latency was <6 msec with the latency jitter <1 msec. Among these, 24% of D1 and 16% of D2 neurons displayed measurable EPSCs upon PCX fiber activation within the OT with 11% of D1 and 14% of D2 neurons receiving monosynaptic excitation (**Fig 5D**). The proportions of D1 and D2 neurons displaying monosynaptic excitation were similar (χ^2^(*1*)=0.009, p=0.923), reflecting that PCX inputs to the OT are not biased towards one cell population. Thus, PCX association fiber activation evokes responses, in some cases monosynaptically, in both D1 and D2 OT MSNs. These results reveal that the PCX can directly influence these two dominant and important striatal neuron populations.

### Topographical organization of PCX neurons innervating OT D1- and D2- type MSNs

Different areas within the anterior PCX may perform unique functions and possess diverse molecular and circuit features (Diodato et al., 2016; Ekstrand et al., 2001; Haberly, 1973; Haberly and Price, 1978; Illig and Haberly, 2003; Large et al., 2017; Luna and Pettit, 2010; Price, 1973). This led us to ask where projections onto OT D1 and D2 MSNs originate from within the anterior PCX? Do OT neurons receive input from a spatially-distributed population of PCX neurons? Or, are the PCX neurons innervating the OT spatially-organized, which would suggest a unique role for PCX cells in that region in the cortico-striatal modulation we uncovered in the previous experiments? To determine the pattern of innervation onto these neuron types, we injected a helper viral vector mixture (AAV9-EF1a-DIO-histone-BFP-TVA and AAV9-EF1a-DIO-RV-G) into the OT of D1-Cre and D2-Cre mice. Two weeks post-infection, a rabies virus (RV-EnvA-ΔG-GFP; (Wickersham et al., 2007)) was injected in the same location. Tissue was then collected one week following for *post-mortem* analyses. Using this model, we subsequently determined the number of rabies virus-labeled (RV+) neurons across: 1) PCX layers, 2) the dorsal/ventral axis, and 3) the rostral/caudal axis. First though, we sought to confirm the glutamatergic identity of the cells innervating the OT, as suggested by the results in **Fig S1D**. While the PCX association fiber network originates from glutamatergic principal neurons (Haberly, 2001; Neville and Haberly, 2004; Shipley and Ennis, 1996), the PCX is comprised of a heterogenous population of both glutamatergic and GABAergic neurons. Therefore, we performed VGlut1 fluorescent *in situ* hybridization on a subset of D1- and D2-Cre mouse tissue following RV infection to confirm that indeed the PCX neurons synapsing upon OT MSNs are glutamatergic (**Fig S6**).

This RV approach successfully infected neurons in the anterior PCX following injection into either D1- or D2-Cre mice, demonstrating that PCX neurons send monosynaptic input to both of these cell populations (**Fig 6A-B**), as also supported by our cell-type specific patch clamp recordings (**Fig 5**). We first quantified the difference in innervation patterns of both cell types across PCX layers. Consistent with our *in vivo* quantification (**Fig 1D**), the majority of neurons innervating both D1 or D2 neurons in the OT originate from within PCX layer ii (D1: *F*(*2,9*)=126.0, *p*<0.0001; D2: *F*(*2,9*)=447.9, *p*<0.0001 comparing across layers; **Fig 6C**). Next, to determine if different PCX subregions uniquely synapse upon these MSN populations, we quantified the number of RV+ cells in each subregion of PCX across D1-Cre and D2-Cre mice. This revealed that more neurons in the ventral versus dorsal PCX synapsed upon these neurons (D1: *t*(*3*)=8.19, *p*=0.004; D2: *t*(*3*)=8.10, p=0.004; **Fig 6D**). Examining the rostral-caudal distribution of PCX innervation of OT we found that more neurons from the caudal versus rostral PCX synapsed upon both D1 and D2 neurons (D1: *t*(*6*)=-12.57, *p*<0.0001; D2: *t*(*6*)=-1.94, *p*=0.01; **Fig 6E**). Thus, a ventral-caudal gradient of PCX neurons innervates, to a similar extent, both OT D1 and D2 MSNs and, therefore, PCX input into the OT is topographically organized.

**Figure 6.**
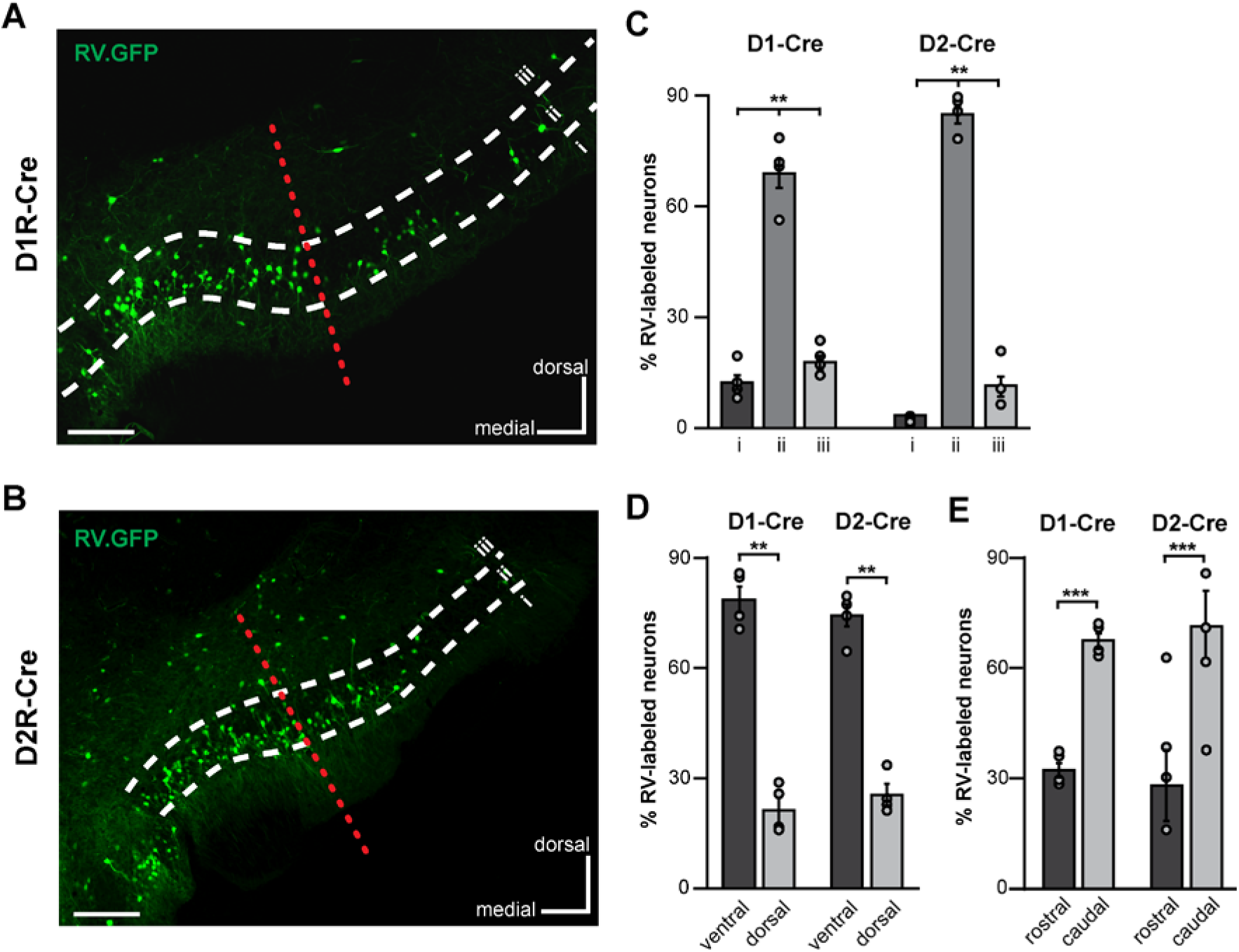
Innervation of OT MSNs by the PCX is topographically organized. **A-B.** Representative coronal brain sections displaying anterior PCX neurons labeled by the rabies virus (RV) (RV-EnvA-ΔG-GFP - green) from the OT in both D1R-Cre (**A**) and D2R-Cre (**B**). Scale bar = 200 μm. **C**. The percentage of RV+ neurons across all PCX layers. Each dot represents the average for one animal. \**p*<0.05, \*\**p*<0.01 for all. **D**. Percentage of RV+ neurons localized within the ventral and dorsal PCX subregions. E. Percentage of RV+ neurons localized within the rostral and caudal PCX subregions. See Fig S6 for confirmation of D1 and D2 MSN innervation by glutamatergic PCX neurons.

## Discussion

Extensive anatomical work over the last century (*e.g.,* (Brunjes et al., 2005; Haberly, 2001; Heimer, 1968; Neville and Haberly, 2004; Scott et al., 1980; Sosulski et al., 2011; White, 1965) has examined the vast and diffuse interconnectivity within the olfactory system. While it is assumed that the way in which the olfactory system disseminates odor information across its expansive hierarchical network is critical for olfactory information processing and ultimately odor perception, the function of this inter-regional connectivity is just beginning to be resolved (Chapuis et al., 2013; Howard et al., 2016; Otazu et al., 2017; Rothermel and Wachowiak, 2014; Sadrian and Wilson, 2015). In the present study we contribute to this overarching goal by defining the function and the cells involved in one specific and unique pathway: the cortico-striatal pathway from the ‘primary’ olfactory cortex into the OT.

### Implications for olfactory processing and perception

We demonstrated that PCX input onto OT neurons bidirectionally modulates an OT neuron’s activity dependent upon the neuron’s intrinsic responsivity to odors. PCX fiber activation decreases the firing of neurons excited by odors, and increases the firing of those suppressed by odors. Thus, the PCX biases OT odor-evoked activity by transforming the population of OT neurons that are odor responsive. This shift of responsivity is likely due to convergent effects of PCX inputs onto certain OT neuron subtypes as we show here, and the simultaneous integration of olfactory bulb input (Schneider and Scott, 1983). This is further made possible given the fact that striatal MSNs extend collaterals upon one another, which while shown to modulate neighboring cells (Taverna et al., 2008), are not well-understood in the context of *in vivo* physiology nor in the context of behavior.

Regarding perceptual and behavioral outcomes of this bidirectional modulation, what would be the significance of exciting neurons which are otherwise suppressed by odor? Similarly, what would be the function of suppressing neurons which are otherwise excited by odor? While the outcome of this upon odor-guided behavior is unknown, we do know that the OT receives monosynaptic odor input from the olfactory bulb in addition to this PCX input we describe herein. So, the OT does not ‘need’ this PCX input to receive likely critical odor information from OB and inform a fundamentally basic odor-guided behavior (‘do I recognize an odor in my environment?’). Instead, the OT likely capitalizes upon the extra richness in odor input from the PCX, including that influenced by perceptual learning and experience (Barnes et al., 2008; Chapuis and Wilson, 2011; Gire et al., 2013; Li et al., 2006). In this model, the OT is now afforded this ‘refined’ odor input due to the bidirectional change upon PCX fiber activation. We predict this refined odor input would facilitate odor-guided behaviors following learning, for instance, in the case of odors with a known reinforced outcome, or valence.

We wish to point out that the amount of OT neuron modulation we observed by PCX activation is likely an underestimation in both our *in vivo* and *in vitro* results. This is especially the case in our paradigms wherein we directly stimulated PCX fibers within the OT itself. First, this may be due to less-than-ideal viral infection efficiency. While a fair number of PCX neurons were infected (**Fig 1**), this number is by far the complete PCX glutamatergic cell population. Second, this could be due to the fact that the AAV injections were not targeted specifically to the ventro-caudal portion of the PCX (which we discovered strongly innervates the OT; **Fig 6**). Further, it is likely we severed connections between these structures during slice preparation or surgical implantation of electrodes and optical fibers. Additionally, it is necessary to consider the direct versus indirect effects of the light stimulation paradigms we used. Stimulation of the PCX locally excites numerous glutamatergic neurons, which extend projections both within the PCX and into additional structures *via* the association fiber network (Haberly, 2001; Neville and Haberly, 2004; Shipley and Ennis, 1996). Thus, changes in OT neuron firing upon local PCX stimulation could be due to a combination of direct PCX inputs and polysynaptic inputs from intermediate structures between the PCX and OT, such as the anterior olfactory nucleus, entorhinal cortex, or amygdala (Haberly, 2001; Neville and Haberly, 2004). Perhaps this is why our direct activation of PCX fibers within the OT yielded a lower impact upon OT firing rates when compared to local PCX activation. This would similarly agree with the smaller population-level monosynaptic light-evoked responses in our *in vitro* preparation. Interestingly, nevertheless, there were significant light-evoked alterations of neuron firing during odor presentation, suggesting that it is a synergistic combination of both odor input and PCX association fiber activation that leads to modulation of OT neuron firing.

Our viral tracing revealed that there is a topographical organization of PCX innervation onto OT D1 and D2 neurons. This is reminiscent of PCX’s topographically organized output upon the orbitofrontal cortex, which also follows along the major axes of the PCX (Chen et al., 2014). Given the distributed and spatially-overlapping representation of odors in the PCX (Poo and Isaacson, 2009; Rennaker et al., 2007; Stettler and Axel, 2009) and other olfactory cortices (*e.g.,* (Kay et al., 2011)), what are the functional implications of this spatially-organized output? The anterior PCX has historically been subdivided into ventral and dorsal regions, with each having differences in access to, and in their responsivity to, sensory input, as well as differences in their expression of molecular markers, and density of cell layers (Diodato et al., 2016; Ekstrand et al., 2001; Haberly, 1973; Haberly and Price, 1978; Illig and Haberly, 2003; Large et al., 2017; Luna and Pettit, 2010; Price, 1973). Further, there is an established rostral-caudal gradient in the magnitude of inhibition onto PCX pyramidal neurons, with inhibition being greater in the caudal PCX (Large et al., 2017; Luna and Pettit, 2010). This effect is largely mediated by layer iii inhibitory neurons, and this inhibitory activity pattern is dependent upon sensory experience (Large et al., 2017). As there is increasing inhibition caudally, and we demonstrated here that more neurons project into the OT from this caudal region, it is possible that these neurons which innervate the OT are subject to differential types of inhibition dependent upon the quality of incoming odor information. Thus, our work adds to other forms of known anatomical and circuit-level heterogeneity within the PCX and therefore our results inform models for how olfactory information is relayed out of the PCX. Additional RV injections into other olfactory structures, and perhaps even into subzones within the OT (the present injections were largely targeted into the antero- medial and antero-lateral OT zones), will be important in defining additional spatial zones within the PCX. It is likely that while the PCX itself represents odors in a distributed, spatially-overlapping manner (Poo and Isaacson, 2009; Rennaker et al., 2007; Stettler and Axel, 2009), the topographical organization of PCX efferents as described herein and in (Chen et al., 2014) allows for the PCX to exert influences which are unique to both the efferent structure, and to odor-evoked activation of select PCX spatial zones.

### The OT as a hub for distributing olfactory input into the basal ganglia

Neurons in the direct pathway, specifically striatonigral MSNs expressing D1 receptors, ultimately disinhibit the thalamus and facilitate motivated behaviors (Gerfen et al., 1990). Neurons of the indirect pathway inhibit the thalamus and diminish motivated behaviors (Calabresi et al., 2014), and are predominantly striatopallidal MSNs expressing the D2 receptor (Gerfen et al., 1990). While we presently do not know specific roles for these cell types within the OT, nor if the strict definition of the direct or indirect pathways applies to these OT cells (Ikemoto and Bonci, 2014; Kupchik et al., 2015; Smith et al., 2013), the OT’s position within the ventral striatum (a component of the basal ganglia) together with our finding that the PCX functionally innervates the OT, adds weight to the possibility that the OT serves as a hub for odor information to enter the basal ganglia. This is further likely since additional olfactory structures also innervate the OT (Wesson and Wilson, 2010). What do our results suggest may subserve this ‘hub’ role for the OT? We propose that our finding that both OT D1 and D2 MSNs equally receive PCX input positions them to serve central roles in this hub – affording the OT to exert and extend odor information into downstream structures which comprise both the direct and direct pathways. By this, PCX input into the OT, or perhaps that arising from another olfactory structure upon the OT, can functionally be distributed into basal ganglia. This routing of odor information into the direct and indirect pathways likely has major implications for odor approach or aversion behaviors. OT D1 MSNs are considered important in mediating the encoding of both appetitive and aversive learned odors with greater numbers than D2 MSNs (Murata et al., 2015). Additionally, dopaminergic input onto OT D1 and D2 MSNs potentiates only D1 MSN responses to olfactory bulb input, suggesting an initial specification of odor information into the OT (Wieland et al., 2015). It is likely that throughout learning, or in states of enhanced motivation, the functional consequence of the observed PCX input may be magnitudes different than that which we report herein. For instance, phasic dopamine release upon OT MSNs may facilitate synaptic input of PCX association fibers and thereby allow for strengthening of PCX modulation. In these cases, the routing of odor information from the OT will be enhanced and thereby so would the robustness of a possible behavioral response (*e.g.,* odor approach). Our results revealing a source of cortical input to this olfactory striatal structure’s MSN population highlights the OT’s role as a possible hub into basal ganglia. We predict that neuromodulators play an integral role in distributing odor information from the OT into other basal ganglia regions.

## Conclusions

We uncovered a functional role for PCX association fibers in exerting modulatory influences upon the OT. The excitatory input upon OT MSN populations by the PCX highlights at least one mechanism whereby the PCX may shape the availability of odor information within an interconnected brain structure. Our finding that this occurred upon both D1 and D2 neuron populations raises the exciting and likely possibility that this pathway informs the display of odor-directed motivated behaviors. As such, one might postulate that the OT, serving as the olfactory striatum and receiving input from numerous olfactory centers (Wesson and Wilson, 2011), participates in concert with other olfactory cortices, beyond the PCX itself, to distribute odor information into critical basal ganglia centers which have direct roles in regulating motor behaviors including stimulus approach and even consummation.

## Methods

### Animals

For *in vivo* experiments, 2-5 months old C57BL/6 male mice (*n*=17 for experiment 1, *n*=23 for experiment 2) originating from Harlan or Jackson Laboratories were bred and maintained within either the Case Western Reserve University School of Medicine or University of Florida animal facility. Mice were housed on a 12 hour light/dark cycle with food and water available *ad libitum* except when water was restricted for behavioral training (see Behavior subsection below). Mice were group-housed up until the point of receiving the intra-cranial implants at which point they were single housed.

For *in vitro* experiments, D1-tdTomato (Shuen et al., 2008) and D2-EGFP (Tg(Drd2-EGFP)S118Gsat) BAC (Gong et al., 2003) transgenic male and female mice (*n*=9; 1-2 months old) were crossed to obtain mice with dopamine D1 and D2 receptor-expressing MSNs labeled in red and green fluorescence, respectively.

All experimental procedures were performed in accordance with the guidelines of the National Institutes of Health and were approved by the Institutional Animal Care and Use Committees at either the University of Florida, Case Western Reserve University, University of Pennsylvania or the Wuhan Institute of Physics and Mathematics, Chinese Academy of Sciences.

### Stereotaxic Surgery and Viral Injections

For *in vivo and in vitro* experiments, mice were anesthetized with isoflurane (2-4% in oxygen; Patterson Veterinary, Greeley, CO) and mounted in a stereotaxic frame with a water-filled heating pad (38°C) to maintain the mouse’s body temperature. Anesthetic depth was verified by the absence of a toe-pinch reflex and respiratory monitoring throughout. A local anesthetic (1% bupivacaine, 0.05 ml, s.c.) injection was administered into the wound margin prior to exposing the dorsal skull. For viral injections, a craniotomy was made above the anterior PCX (A/P: −1.0mm, M/L: +2.8mm, D/V: +3.5mm), and either a 33Ga Hamilton microsyringe or a glass micropipette was lowered into the PCX. 0.5 μl of AAV5.CaMKiiα.hChR2.E123T.T159C.p2A.mCherry.WPRE (cell-filling variant) or AAV5.CaMKiiα.hChR2(H134R).mCherry (non-cell-filling variant), or control vector AAV5.CaMKiiα.mCherry.WPRE (all undiluted, University of North Carolina Viral Vector Core, Chapel Hill, NC, USA), was infused by a pump at a rate of 0.05 μl/min. The syringe/pipette was gently withdrawn, the craniotomy sealed with wax, and the wound margin closed. During the recovery period, mice received a daily injection of carprofen (Pfizer Animal Health) or meloxicam (Patterson Veterinary; 5 mg/kg, s.c. for both) and allowed *ad libitum* access to food and water.

For *in vivo* experiments, after allowing 2-3 weeks for the viral vector to transduce in the PCX, we again performed stereotaxic surgery to either implant an optical fiber in the PCX and electrode array in the OT (for PCX stimulation), or an optetrode into the OT (for PCX association fiber stimulation). For fiber and array implants, a craniotomy was made dorsal to the anterior PCX and a glass optical fiber was lowered into the PCX. Another craniotomy was made dorsal to the OT, and an 8-channel tungsten wire electrode was lowered into the OT. For optetrode implants, a craniotomy was made dorsal to the OT, and an optetrode assembly was lowered ~0.3 mm dorsal to the OT. All craniotomy sites were sealed with wax and the implants cemented in place, along with a head bar for head-fixation during behavioral experiments (see Behavior subsection below). Mice recovered for 5 days prior to beginning behavioral experiments, and received a daily injection of carprofen or meloxicam (same doses as above) and *ad libitum* access to food and water during this time.

For viral tracing experiments, viral tools for trans-mono-synaptic labeling were packaged by BrainVTA (BrainVTA Co., Ltd., Wuhan, China). The helper virus AAV9-EF1a-DIO-histone-BFP-TVA and AAV9-EF1a-DIO-RV-G were titrated at about 3×1012 genome copies per milliliter, and RV-EnvA-ΔG-GFP was titrated at about 2x108 infecting unit per milliliter. The mixture of AAV9-EF1a-DIO-histone-BFP-TVA and AAV9-EF1a-DIO-RV-G (volume ratio: 1:1, 100 nl in total) was injected into the OT (A/P: +1.2mm, M/L: +1.1mm, D/V: +5.5mm) in D1 R-Cre (*n*=4) and D2R-Cre (*n*=4) mice, respectively. Two weeks later, 150nl of RV-EnvA-ΔG-dsRed was injected into the same location of these mice. One week after the RV injection, the mice were perfused transcardially with PBS (Pre-treated with 0.1% diethylpyrocarbonate (DEPC, Sigma)), followed by ice-cold 4% paraformaldehyde (PFA, 158127 MSDS, Sigma)

### Behavior and Stimulus Presentation

Following surgical recovery, mice for *in vivo* experiments were water-restricted on a 23-hour schedule to no less than 85% of their body weight to induce motivation to perform the task. For the fixed-interval olfactory task, mice were head-fixed in a tube with an olfactometer and lick spout directly in front of the mouse’s snout and mouth, respectively. Mice were first habituated to head-fixation across 2-3 days prior to stimulus presentation and were head-fixed for no more than 15 minutes per session. During acclimation, mice were lightly anesthetized with isoflurane prior to head fixation to minimize stress. Mice then began learning the fixed-interval olfactory task, with the following structure for each trial: 1) trial start for 12 seconds, 2) 1 of 9 pseudorandomly presented stimuli for 2 seconds; 3) 8 second post-stimulus rest period; 4) access to fluid reinforcer (2 mM saccharin) for 10 seconds prior to new trial start. The timing of the stimulus and reward presentation ensured that there were no lick movement artifacts in the multi-unit recordings due to anticipation of reward, and that all stimuli were equally rewarded. Thus, across a session and over days, mice learned to lick for reward to all stimuli presented and worked up to at least 9 trials of each stimulus presentation (~1 hour session/day). Pseudorandomly-presented stimuli were one of the nine following: 4 monomolecular odors (isopentyl acetate, heptanal, ethyl butyrate, 1,7 octadiene; Sigma-Aldrich, St. Louis, MO), the same 4 odors + 10-Hz (20 ms pulse width) PCX light stimulation, or light stimulation alone. Odors were delivered via an air-dilution olfactometer via independent lines of tubing. All odors were diluted to 1 Torr in mineral oil and delivered to the olfactometer via medial-grade nitrogen at a flow rate of 1L/min. Mice needed to lick for the fluid reinforcer post-stimulus in at least 85% of trials for the session to be included for analysis of neural activity. Trials were divided into correct (licked for fluid reinforcer after stimulus) and incorrect (did not lick for reinforcer after stimulus presentation) responses.

### Electrophysiology and Optical Stimulation

#### Probe fabrication

For *in vivo* experiments, all optical fibers, electrode arrays, and optetrodes were custom-made. For optical fibers, glass multimode fiber (300 μm core, 0.39NA; Thorlabs, Newton, NJ) was cut to the appropriate length and fastened with optical adhesive (Norland Products, Cranbury, NJ) in a 2.5 mm ceramic ferrule (Thorlabs). Fiber ends were smoothed with lapping pads for ceramic ferrules (Thorlabs). For fixed electrode arrays, tungsten wire (A-M Systems, Sequim, WA) was attached to an Omnetics connector via silver epoxy, with a stainless steel wire (A-M Systems) serving as the ground wire. Tungsten wires were bundled in two polyimide tubes and cut to the appropriate length. Optetrode assembly was performed in accordance with (Anikeeva et al., 2011), with only slight modifications made to reach the appropriate ventral depth. Light intensity output of the final pre-implanted fiber was 7-10 mW^3^ and the distance between tetrode tips and fiber end were 500-800 μm apart to yield a broad light cone for neuron activation.

#### In vivo electrophysiology and optical stimulation

The connector of the fixed electrode array or optetrode was connected to a headstage and electrode channels were digitized to acquire multi-unit activity (24 kHz, 200-3kHz band-pass filter) and lick events (300 Hz) for behavioral criterion monitoring. Light stimulation was provided by 447.5nm LED (Luxeon Rebel ES, Luxeon Stars, Lethbridge, Alberta) driven by a Thorlabs T-Cube LED driver (Thorlabs) connected to a fiber patch cable (ThorLabs; 300μm core multimode fiber) for temporary connection to the implanted ferrule on the animal. Light stimulus intensity, timing, and frequency were determined after conducting pilot experiments where we stimulated the PCX between 5-30 Hz at 20 ms pulse width, and intensity was adjusted for every mouse to the lowest intensity possible while still evoking light-induced responses in the OT. For PCX association fiber stimulation experiments, tetrodes on the optetrode were advanced 50μm/session to access new neurons within the OT. For the fixed array, 1 of the 8 channels from the multiunit recordings was used as a local reference channel for signal subtraction.

#### In vitro electrophysiology and optical stimulation

For *in vitro* whole-cell patch clamp recordings, mice were deeply anesthetized with ketamine-xylazine (200 and 20 mg/kg body weight, respectively) and decapitated. The brains were dissected out and immediately placed in ice-cold cutting solution containing (in mM) 92 N-Methyl D-glucamine, 2.5 KCl, 1.2 NaH2PO4, 30 NaHCO3, 20 HEPES, 25 glucose, 5 Sodium L-ascorbate, 2 Thiourea, 3 Sodium Pyruvate, 10 MgSO4, and 0.5 CaCl2; osmolality ~300 mOsm and pH ~7.3, bubbled with 95% O2-5% CO2. Coronal sections (250 μm thick) containing the PCX and OT were cut using a Leica VT 1200S vibratome. Brain slices were incubated in oxygenated artificial cerebrospinal fluid (ACSF in mM: 124 NaCl, 3 KCl, 1.3 MgSO4, 2 CaCl2, 26 NaHCO3, 1.25 NaH2PO4, 5.5 glucose, and 4.47 sucrose; osmolality ~305 mOsm and pH ~7.3, bubbled with 95% O2-5% CO2) for ~30 min at 31 °C and at least 30 minutes at room temperature before use. For recordings, slices were transferred to a recording chamber and continuously perfused with oxygenated ACSF. Fluorescent cells were visualized through a 40X water-immersion objective on an Olympus BX61WI upright microscope equipped with epifluorescence.

Whole-cell patch clamp recordings were made under both current and voltage clamp mode. Recording pipettes were made from borosilicate glass with a Flaming-Brown puller (P-97, Sutter Instruments; tip resistance 5-8 MΩ). The pipette solution contained (in mM) 120 K-gluconate, 10 NaCl, 1 CaCl2, 10 EGTA, 10 HEPES, 5 Mg-ATP, 0.5 Na-GTP, and 10 phosphocreatine. Electrophysiological recordings were controlled by an EPC-9 amplifier combined with Pulse Software (HEKA Electronic) and analyzed using Igor Pro. The signals were filtered at 2.9 kHz and acquired at 50 kHz. Excitatory postsynaptic potentials (EPSPs) were further filtered offline at 20 kHz and excitatory postsynaptic currents (EPSCs) at 0.5 kHz. Junction potential (~9 mV) was corrected offline. Light stimulation was delivered through the same objective via pulses of blue laser (473 nm, FTEC2473-V65YF0, Blue Sky Research, Milpitas, USA) with varying lengths. Viral infection in the PCX was confirmed in brain slices during recordings. No sex differences between male and female mice were evident so data was pooled across mice.

### Histology

Histological confirmation of PCX and OT implant sites for *in vivo* experiments was performed using a DAPI (4’,6-diamidino-2-phenylindole, Invitrogen, Carlsbad, CA; **Fig S2Ai-ii**) stain of 40 μm coronal brain sections. The number of cells transduced in the PCX was quantified in 4-6 alternate 20 μm sections within a set region of interest in a subset of animals (300μm DV × 500μm ML/section; *n*=4), and compared to the number of anti-NeuN-labeled neurons to provide an estimate of AAV transduction using Nikon NIS Elements. The spread of AAV infection was also estimated across all experimental animals using Nikon NIS Elements software (*n*=12 mice, 0.87mm ± 0.04mm anteriorposterior spread relative to bregma, see **Fig S2C**). For VGlut1 confirmation, the number of PCX cells coexpressing viral transduction and VGlut1 was examined in 4-6 alternate 40 μm sections in a subset of mice (*n*=3; **Fig S2D**). For anti-NeuN and VGlut1 immunohistochemistry, free-floating sections were rinsed in tris-buffered saline and diluting buffer, and then blocked in 20% normal donkey serum for 30 minutes. Slices were then incubated for 24 hours at 4°C with the primary antibody anti-VGlut1 antibody (anti-VGlut1: 1:500 in diluting buffer, Millipore, Burlington, MA; anti-NeuN: 1:1000 in diluting buffer, Abcam, Cambridge, UK). Sections were rinsed with diluting buffer and then incubated in with secondary antibody (anti-VGlut1: donkey anti-guinea pig IgG, FITC conjugate, 1:375 in diluting buffer, Millipore; anti-NeuN: donkey anti-rabbit IgG, AlexaFluor 488, 1:500 in diluting buffer, Abcam) for 2 hours at room temperature before being rinsed with tris-buffered saline and then dH20. Tissue was then mounted on slides with DAPI (for anti-VGlut1) or Vectashield mounting medium (for anti-NeuN, VectorLabs, Burlingame, CA) and imaged. To assess association fiber density in the OT, the fluorescent intensity of each layer was calculated as a function of background intensity in NIH ImageJ (Schneider et al., 2012) in 5-6 alternate 40 μm sections in a subset of mice (*n*=4). Data from animals with either unrestricted or unsuccessful labeling were excluded from analysis.

To determine the distribution of RV+ cells in the anterior PCX for viral tracing experiments, 40 μm coronal sections were stored at −20°C floating in 20% glycerine, 30% glycol in PBS. A subset of coronal sections across the anterior PCX (from ~2.34mm – 0.14mm bregma, ~ every 240 μm) were washed in PBS and then mounted onto Superfrost Plus slides with 90% glycerol in PBS and sealed with nail polish. Images of these sections were captured with the Olympus VS120 virtual microscopy slide scanning system (Olympus, Shanghai, China) using a 10x objective. PCX layers, borders, and rostral-caudal axis were delineated based on a standard brain atlas (Paxinos and Franklin, 2000). The dorsal and ventral subregions of the anterior PCX were divided based on the dorsal edge of the lateral olfactory tract (Ekstrand et al., 2001).

To confirm that OT neurons are innervated by glutamatergic neurons in the anterior PCX, we performed fluorescent in situ hybridization (FISH) using a VGlut1 probe on the rabies-labeled samples. Anterior PCX coronal sections (15 μm thick) were collected on Superfrost Plus slides and stored at −80°C. Sections were blocked with 1% hydrogen peroxide in PBS for 15 min at room temperature, rinsed with PBS 2x5 min, and incubated with 1μg/ml Proteinase K (Sigma, P-6556) for 10 min at 37°C. After a 0.2% glycine in PBS rinse for 3 min, sections were then incubated with 0.25% acetic anhydride in 0.1 M triethanolamine, pH 8.0, for 15 min at room temperature and washed again with PBS. VGlut1 probes were pre-diluted to 0.2ng/μl with the hybridization buffer (50% formamide, 300μg/ml tRNA, 10 mM Tris pH 7.5, 10% dextran sulfate, 1x Denhalt’s solution, 600mM NaCl, 0.25% SDS), mixed well, preheated at 80°C for 10 min, cooled with ice, and applied to each section. After overnight incubation at 56°C, the sections were washed, first with 2x SSC for 5 min, and then 0.2x SSC twice for 20 min at 65°C. Once cooled to room temperature, sections were incubated with peroxidase-conjugated anti-DIG antibody (1:500, Roche Applied Science) at 37°C for 1 hour, washed with PBST for 3x5 min, and lastly trated using TSA-plus Cy3 kit (Perkin Elmer). Since the GFP signal from RV labeled called was quenched during the ISH process, we further performed the immunohistochemical staining for GFP with rabbit anti-GFP (1:1000, Abcam) and treated the sections with Alexa 488-conjugated goat anti-rabbit secondary (1:1000, Jackson ImmunoResearch), and DAPI and then mounted and sealed as described above. Images were captured as designated above. RV labeled neurons in different anterior PCX subregions and VGlut1 colabeled neurons in the anterior PCX were quantified semi-automatically using FIJI with the cell counter plugin.

### Data Analysis

Behavioral and electrophysiological data analyses were performed using custom scripts (Cambridge Electronic Design, MATLAB). Neurons were sorted offline in Spike2 using template matching and principal component analysis, and spike times exported to MATLAB for analysis. Any putative single unit with >2% of events occurring within a 2 msec inter-spike interval were excluded from analysis (Gadziola et al., 2015). Significance level for all statistical tests was set at *p*<0.05, and all *t*-tests performed were paired unless otherwise noted. Significant light-evoked units were defined by comparing the average light-evoked response across 2sec of light stimulation to 2sec of spontaneous activity (prior to light onset). Odor-excited units are defined as those units whose firing rates were significantly increased during the odor presentation period (2sec) when compared to spontaneous firing rates (2sec prior to odor onset). Odor-suppressed units significantly decreased their firing rates during odor presentation (2sec) when compared to spontaneous firing rates (2sec prior to odor onset). Significant changes in the number of odor-responsive neurons with simultaneous light stimulation were determined by using a χ^2^ with Yates’ correction. A Komolgorov-Smirnoff test was used to examine the population-level change in the magnitude of firing rates from odor to odor+light. A one-sample z-test for proportions was used to quantify the differences between the proportion of neurons that were RV+ and VGlut1+ versus those that were only RV+.

## Author Contributions

Conceptualization: K.A.W. and D.W.W.; Methodology: K.A.W., Z.Z., F.X., M.M., and D.W.W.; Investigation: K.A.W., Y-F.Z., Z.Z., J.P.B., A.H.M., E.I.Z., H.M., and X.J.; Resources: M.V.F.; Writing – Original Draft: K.A.W., Z.Z., M.M., and D.W.W.; Writing – Review & Editing: all authors; Visualization: K.A.W., Z.Z., Y-F.Z., J.P.B., D.W.W.; Supervision: F.X., M.M., and D.W.W.; Funding Acquisition: K.A.W., F.X., M.M., and D.W.W.

## Acknowledgements

This work was supported by NIH NIDCD R01DC014443 and R01DC016519 to D.W., R01DC006213 to M.M., F31DC016202 to K.W, and National Natural Science Foundation of China grant 31771156 to F.X. We thank Marie Gadziola for help with *in vivo* data handling and Chris Ford for advice on medium spiny neuron recordings. We thank Zhonghua Lu and Liping Wang from the Shenzhen Institutes of Advanced Technology for providing the VGlut1 probe and platforms for ISH, Yanqiu Li from the Wuhan Institute of Physics and Mathematics (WIPM) for providing the D1R-Cre/D2R-Cre mice used in viral tracing, and Lingling Xu from WIPM for managing the microscopy platform.

